# Determinants for forming a supramolecular myelin-like proteolipid lattice

**DOI:** 10.1101/2020.02.06.937177

**Authors:** Salla Ruskamo, Oda C. Krokengen, Julia Kowal, Tuomo Nieminen, Mari Lehtimäki, Arne Raasakka, Venkata P. Dandey, Ilpo Vattulainen, Henning Stahlberg, Petri Kursula

**Author notes:** equal contribution (Salla Ruskamo, Oda C. Krokengen). J.K.; Institute of Molecular Biology and Biophysics, ETH Zurich, Switzerland., T.N.; Tampere University of Applied Sciences, Tampere, Finland, M.L.; FDA, Rockville, MD, USA, V.P.D.; The National Resource for Automated Molecular Microscopy, Simons Electron Microscopy Center, New York Structural Biology Center, New York, NY, USA.

## Abstract

Myelin protein P2 is a peripheral membrane protein of the fatty acid binding protein family. It functions in the formation and maintenance of the peripheral nerve myelin sheath, and several P2 mutations causing human Charot-Marie-Tooth neuropathy have been reported. Here, electron cryomicroscopy of myelin-like proteolipid multilayers revealed a three-dimensionally ordered lattice of P2 molecules between stacked lipid bilayers, visualizing its possible assembly at the myelin major dense line. A single layer of P2 is inserted between two bilayers in a tight intermembrane space of ∼3 nm, implying direct interactions between P2 and two membrane surfaces. Further details on lateral protein organization were revealed through X-ray diffraction from bicelles stacked by P2. Surface mutagenesis of P2 coupled to structural and functional experiments revealed a role for both the portal region and the opposite face of P2 in membrane interactions. Atomistic molecular dynamics simulations of P2 on myelin-like and model membrane surfaces suggested that Arg88 is an important residue for P2-membrane interactions, in addition to the helical lid domain on the opposite face of the molecule. Negatively charged myelin lipid headgroups anchor P2 stably on the bilayer surface. Membrane binding may be accompanied by opening of the P2 β barrel structure and ligand exchange with the apposing lipid bilayer. Our results provide an unprecedented view into an ordered, multilayered biomolecular membrane system induced by the presence of a peripheral membrane protein from human myelin. This is an important step towards deciphering the 3-dimensional assembly of a mature myelin sheath at the molecular level.

## Introduction

A central question in myelin biology is the molecular mechanism of the tight packing of dozens of apposing lipid bilayers into a mature, multilayered myelin sheath. A major role in this process is played by myelin-specific proteins. The high degree of order within the myelin sheath has been known since early experiments using X-ray diffraction ^1^; however, the details of the molecular assembly have remained enigmatic.

The spontaneous formation of lipid membrane multilayers is a common functional property of different myelin-specific proteins, which are not genetically related. In peripheral nervous system (PNS) myelin, the compact multilamellar membrane contains only a few proteins. The intrinsically disordered myelin basic protein (MBP) is irreversibly embedded into a single leaflet of the lipid bilayer ^2^. The cytoplasmic domain of myelin protein zero (P0) behaves much like MBP, although it embeds deeper into the membrane ^3^. Full-length P0 promotes membrane stacking through both extra- and intracellular interactions ^3–6^. Peripheral myelin protein 22 (PMP22), another PNS integral membrane protein, forms myelin-like assemblies ^7^, much like those observed with MBP and P0. P2 adheres to the cytoplasmic leaflet of the bilayer and can be classified as a peripheral membrane protein ^8^.

Peripheral membrane proteins associate with cellular membranes *via* diverse mechanisms. Membrane binding may be either irreversible, mediated by post-translational modifications (palmitoylation, myristoylation, or prenylation), or reversible with variable binding affinities. The specificity of protein-membrane interactions is affected by the physical properties of the protein and the lipid bilayer, such as surface charge or membrane curvature. Many peripheral membrane proteins utilize amphipathic helices or hydrophobic amino acids that penetrate into the hydrophobic bilayer core to form stable interactions with membranes ^9^.

P2 is a Schwann cell-specific protein expressed in the PNS myelin of tetrapods ^10^. Intriguingly, P2 is expressed in a mosaic fashion, not being present in all myelin sheaths ^11,12^. This small β-barrel protein belongs to the family of fatty acid binding proteins (FABPs). The bound fatty acid is enclosed inside the β barrel by a lid formed by two adjacent α helices ^13–15^; the opening of the β barrel may be of importance in fatty acid entry and egress ^13^. In addition to fatty acid binding, P2 can transfer lipids from/to membranes using a collisional transfer mechanism ^16^, as seen with several other FABPs ^17–21^. Besides fatty acids, P2 may bind cholesterol ^14^, which is abundant in the myelin membrane and essential for myelination ^22^. The tip of the α-helical lid is hydrophobic, while both ends of the β barrel present positively charged surfaces ^14,15^, and these properties are likely important, when P2 stacks between two phospholipid bilayers.

Studies on P2-deficient mice revealed temporarily reduced motor nerve conduction velocity and altered lipid composition in PNS myelin. However, the overall PNS myelin structure remained normal ^16^. Further analyses on the mutant mice revealed that P2 has a role in remyelination of an injured PNS ^23^ and melanoma cell invasion ^24^. Five Charcot-Marie-Tooth 1 (CMT1) disease point mutations in human P2 have been discovered ^25–29^. Three CMT1-associated P2 protein variants have been characterized at the molecular level, showing altered fatty acid and lipid membrane binding properties. The most drastic CMT1 mutation, T51P, also reduced the membrane stacking capability of P2 ^30^. Overall, the stability of the mutant proteins was decreased, even though crystal structures indicated only minor structural changes compared to wild-type P2.

In the current study, we incorporated human P2 into a model membrane multilayer system and visualized the myelin-like proteolipid structures using electron cryomicroscopy (cryo-EM). P2-bicelle complexes were used for additional structural insights. We produced mutated forms of P2 to establish determinants of lipid bilayer and fatty acid binding and used atomistic molecular dynamics (MD) simulations to visualize the intimate interaction between P2 and a myelin bilayer. We show the spontaneous formation of an ordered, crystal-like lattice of P2 bound inside membrane multilayers and highlight factors that are important in this process, which involves a conformational change in the protein. The results provide a glimpse into the self-assembling properties of myelin proteins and lipid membranes, which are likely to be crucial for correct myelination in the vertebrate nervous system.

## Materials and methods

### Protein production

The expression and purification of human wild-type P2 (wtP2) was done as described ^14^. Mutagenesis and the expression and purification of P2 variants were described earlier ^31^.

### Electron cryomicroscopy and image processing

0.6 mg/ml of purified wtP2 or the P38G variant were mixed with *E. coli* polar lipids (Avanti Polar Lipids) using a lipid:protein ratio of 2 (w/w), corresponding to a molar ratio of ∼40, and incubated for 1-2 h at +23 °C. For grid preparation, samples were applied to glow-discharged, holey carbon grids (QUANTIFOIL R 1.2/1.3, R 2/2 or R 3.5/1). 3-µl samples were adsorbed for 1 min at +20 °C, 90% humidity. Grids were then blotted for 2 s and vitrified by plunging into liquid nitrogen-cooled liquid ethane using an FEI Vitrobot MK4 (Vitrobot, Maastricht Instruments). The frozen grids were imaged using FEI Titan Krios TEM operated at 300 keV. Images were recorded using a Gatan K2 Summit direct electron detector, in counting mode (0.2 sec/frame, 8 sec in total, 6-7 e/pix/sec). Movie frames were aligned with MotionCorr ^32^ and preprocessed by 2dx_automator ^33^. The effective pixel size of the images was 1.3 Å/pixel. Particles were boxed with EMAN2 (Helixboxer) ^34^ and further processed by Spring ^35^ with helical reconstruction. In total, 25 000 overlapping and CTF-corrected segments with the size of 240 × 240 pixels were used with a binning value of 2 to calculate 2D class averages.

### Structural analysis of P2-stacked bicelles

0.5 mg/ml P2 was mixed with 0.5 mg/ml bicelles (phospholipid:dodecylphosphocholine (DPC) ratio 2.85, phospholipids 1:1 dimyristoylphosphatidylcholine (DMPC):dimyristoylphosphatidylglycerol (DMPG) and incubated for 1 h at room temperature. 4-µl samples were then pipetted onto glow-discharged carbon-coated copper grids before incubating for 1 min. Excess solution was removed with filter paper (Whatman), and the samples were washed with 4 drops of Milli-Q water. Samples were stained with two drops of 2% uranyl acetate for 12 s in each drop and air-dried. Transmission electron microscopy (TEM) was performed using a Jeol JEM-1230 (MedWOW) instrument.

To examine repetitive structures in turbid samples, 2, 10, and 20 µM P2 was mixed with 1, 2, or 3 mM bicelles in 20 mM HEPES (pH 7.5), 150 mM NaCl. Samples were prepared at ambient temperature right before the measurements and measured at +25 °C. Synchrotron SAXS data from the suspensions were collected at the PETRA III storage ring, DESY, Hamburg, Germany on the EMBL beamline P12 ^36^. Data were processed and analysed using ATSAS ^37^. Repeat distances in the sample were deduced from Bragg peak positions.

### Crystal structure determination

All P2 variants were crystallized and X-ray diffraction data collected as described ^31^. Data were processed with XDS ^38,39^, and molecular replacement was done using Phaser ^40^ using human wtP2 (PDB code 2WUT) ^14^ as a search model. Structures were refined with phenix.refine ^41^, and rebuilding was done in Coot ^42^. The structures were validated using MolProbity ^43^. The refined coordinates and structure factors were deposited at the PDB (see Supplementary Table 1 for statistics and entry codes).

### Proteolipid vesicle aggregation

5 µM of each P2 mutant was mixed with DMPC:DMPG (1:1) vesicles in a buffer containing 10 mM HEPES (pH 7.4), 150 mM NaCl and incubated for 10 min at room temperature. Lysozyme and BSA were used as negative controls. Turbidity was measured on a Tecan Infinite M200 plate reader at 450 nm. The turbidity values were plotted as relative turbidity compared to wtP2 from the same measurement series. For further characterization, the turbid samples were centrifuged and the supernatant and pellet fractions analyzed by SDS-PAGE, in order to detect cosedimentation of P2 with aggregated vesicles.

Turbidity was also studied with protein-bicelle complexes, using wtP2. For this purpose, bicelles (phospholipid/detergent ratio 2.85) were prepared as above, but the phospholipid composition was varied. Bicelles were at 5 mM and P2 at 33 µM. wtP2 was simultaneously used to examine the effect of phospholipid vesicle composition on protein-induced turbidity; the lipid concentration was 0.5 mM. Turbidity was measured using a Tecan Spark 20M microplate reader at +30 °C.

### Surface plasmon resonance

Surface plasmon resonance (SPR) was used to determine the effect of mutations on P2 to the binding of the protein to lipid monolayers using the Biacore T100 SPR instrument. Lipid monolayers consisting of either DMPC or dimyristoyl phospatidic acid (DMPA) were immobilized on an HPA chip (GE Heathcare) according to the manufacturer’s instructions. P2 at 1.0 μM was injected onto the chip at +25 °C using 10 mM HEPES (pH 7.4), 150 mM NaCl as running buffer.

### Circular dichroism spectroscopy

Circular dichroism (CD) spectra were measured in 10 mM sodium phosphate (pH 7.0) at a protein concentration of 0.1 mg/ml, using quartz cuvettes with 0.1-cm pathlength and a Jasco J-715 spectropolarimeter. Melting curves were measured using 0.2 mg/ml protein at 217 nm. The temperature was increased 1 °C/min from +20 °C to +90 °C.

Synchrotron radiation CD (SRCD) measurements for selected variants were performed on the CD1 beamline of the ASTRID storage ring at the ISA synchrotron (Aarhus, Denmark). Scans from 280 to 165 nm were performed in 1-nm steps at +20 °C in H_2_O. Three mutants with large effects on membrane binding (L27D, R30Q, and P38G), were studied in a bicelle environment (4:1 DMPC/DPC).

Bicelles and vesicles with varying phospholipid compositions were used for further SRCD experiments with wtP2. 0.4 mg/ml wtP2 was mixed with 5 mM bicelles or 2.65 mM vesicles. Spectra were recorded from 280 to 170 nm at +30 °C, using a 100-µm cuvette. These experiments were performed on the AU-CD beamline of the ASTRID2 storage ring at the ISA synchrotron (Aarhus, Denmark)

### Fluorescence spectroscopy

The fluorescent fatty acid 11-dansylamino-undecanoic acid (DAUDA) was used to study fatty acid binding by P2. DAUDA has been used to study ligand binding in FABPs before ^44,45^. DAUDA was dissolved in DMSO, and the final DMSO concentration in the samples was 1%. 20 µM DAUDA was mixed with 0, 1, 5, or 10 µM protein. Samples were incubated for 2 h at +23 °C. Fluorescence excitation at 280 nm was used and emission was recorded at 530 nm using a Tecan Infinite M200 plate reader.

The binding of wtP2 and the P38G mutant to cholesterol was studied using the environment-sensitive fluorescent cholesterol analogue 22-(N-(7-nitrobenz-2-oxa-1,3-diazol-4-yl)amino)-23,24-bisnor-5-cholen-3β-ol (22-NBD-cholesterol). The fluorescence intensity of 22-NBD-cholesterol increases and the fluorescence emission maximum shifts, if the probe is moved to a non-polar environment. 100 µM 22-NBD-cholesterol stock solution was prepared in 100% ethanol, and the maximum ethanol concentration in the sample was 2%. All experiments were carried out in 10 mM HEPES (pH 7.5). 2 µM 22-NBD-cholesterol was incubated for 16 h at +23 °C with varying amounts of P2. Fluorescence spectra were recorded on a Horiba Fluoromax-4 instrument, using excitation at 473 nm and emission between 500-600 nm, with a bandwidth of 5 nm.

### Atomic scale molecular dynamics simulations

Structures of wtP2 and P38G were prepared for the simulations essentially as described elsewhere ^46^. Briefly, the P2 structure with bound palmitate was taken from the PDB entry 4BVM ^15^ and converted to match an all-atom representation consistent with the CHARMM36 force field ^47^, which was used for simulating the system components, unless mentioned otherwise. The topology for wtP2 was directly obtained from this conversion. The P38G mutation was made *in silico* and equilibrated in a water environment. Both protein-palmitate complexes had a total charge of +10.

Lipid bilayers were constructed using the CHARMM-GUI membrane builder ^48^. Two different membrane systems were considered: a 1:1 DMPC:DMPG bilayer as a general reference with a net negative surface charge, and a myelin bilayer mimicking the cytoplasmic leaflet of the myelin membrane. The composition of the myelin-like bilayer was 44 mol-% cholesterol, 27 mol-% palmitoyloleoylphosphatidylethanolamine (POPE), 2 mol-% phosphatidylinositol-4,5-bisphosphate (PIP_2_), 11 mol-% palmitoyloleoylphosphatidylcholine (POPC), 13 mol-% palmitoyloleoylphosphatidylserine (POPS), and 3 mol-% sphingomyelin ^49^. The bilayers were symmetrical, comprised of a total of 200 lipid molecules each.

Ten Cl^-^ ions were included to neutralize the total charge of each protein-palmitate complex. The systems were solvated with a total of 15000 water molecules each, with 0.1 M KCl. Water was modelled using the TIP3P model ^50^. Additional counterions (90 K^+^ in the DMPC:DMPG and 32 K^+^ in the myelin membrane system) were included to neutralize the system total charge. The total system volume was approximately (7.5 × 7.5 × 12) nm^3^ for the DMPC:DMPG and (6.5 × 6.5 × 13.5) nm^3^ for the myelin membrane systems. Periodic boundary conditions were used to make the bilayer structure continuous.

Molecular dynamics (MD) simulations were carried out under NpT conditions. Temperature coupling was performed with the velocity-rescale method ^51^, using separate thermostats for the protein, the bilayer, and the solvent. Reference temperatures were set at 310 K, with coupling time constants of 2.0 ps. Pressure coupling was done semi-isotropically with the Parrinello-Rahman barostat ^52^, using reference pressures of 1.0 bar with coupling time constants of 2.0 ps and compressibility constants of 4.5 × 10^−5^ bar^-1^.

All bonds were constrained with the LINCS algorithm ^53^. Cut-off radii of 1.0 nm were introduced for the Coulombic and Lennard-Jones interactions, including the neighbour list. Long-range electrostatics were calculated using the particle-mesh Ewald method ^54^ with cubic interpolation and a spacing of 0.16 nm for the Fourier grid.

The simulation systems were built by adding the protein structure near the bilayer and solvating the system thereafter. After a short steepest-descent equilibration, the systems were simulated long enough for the protein to spontaneously come into close contact with the bilayer. This was used as the starting structure, after which the systems were simulated for 3 µs each. A total of 4 full simulations were run (one for each protein-membrane combination), in addition to several shorter simulations on the P2 membrane attachment phase. The first 500 ns of each simulation were removed as an equilibration period, and the final 2.5 µs were used for analyses. All simulations were conducted with GROMACS 4.6.7 ^55^, using the CHARMM36 all-atom representation and a time step of 2 fs, saving the trajectory coordinates every 50 ps.

## Results

While the molecular composition of compact myelin is relatively simple, the arrangement of proteins within the membrane multilayers is to a large extent unknown. Here, we used the peripheral membrane protein P2 from the PNS myelin major dense line as a model system to study myelin-like membrane stack formation and structure. P2 interacts with lipid bilayers with high affinity ^13,15,56–58^. We explored its membrane binding characteristics, determinants, and dynamics more closely. The results provide further information on the molecular details of the major dense line in PNS myelin, as well as on CMT disease mechanisms linked to mutations in P2.

### Arrangement of P2 in multilayered membrane stacks

P2 spontaneously binds lipid membranes together, as reflected by earlier studies using turbidimetry, simulation, and X-ray diffraction ^15,57^. However, the molecular details of this phenomenon and the resulting supramolecular structure have remained elusive. Cryo-EM was used to follow membrane stacking and ordering of proteolipid components in multilayers induced by P2.

P2 induced the formation of highly ordered lipid bilayer stacks, while without P2, only unilamellar vesicles were observed (Figure 1A,B). The angle between two separating bilayers at the edge of a tight apposition is consistently <60 ° (Figure 1C). Although P2 is only 15 kDa, it is visible in cryo-EM images as ordered rows of particles between two apposed membranes. Based on the calculated 2D class averages (Figure 1D-G), P2 evidently stabilizes the lipid membrane stacks and defines the spacing (∼3.0 nm) between two bilayer surfaces. This myelin-like spacing between two apposing lipid bilayers is constant throughout the membrane stacks. Based on the P2 crystal structure ^14,15^, the longest diameter of P2 is 4.5 nm, indicating that either some parts of the protein are buried within the bilayer, or P2 is turned on its side on the membrane. The repeat distance in the multilayer, containing a 4.5-nm bilayer and a single layer of P2 molecules, is 7.5 nm. This is shorter than the distance measured in solution with X-ray diffraction under more hydrated conditions, and close to the distance observed with MBP and the P0 cytoplasmic domain in diffraction experiments ^2,3,15^. P2 molecules are located between the bilayers with a lateral spacing of 3.5 nm between monomers (Figure 1F), indicating lattice-like order between membranes. This order extends into neighbouring membrane layers, and P2 molecules between the bilayers are at least to some extent in register between consecutive layers (Figure 1G).

**Figure 1.**
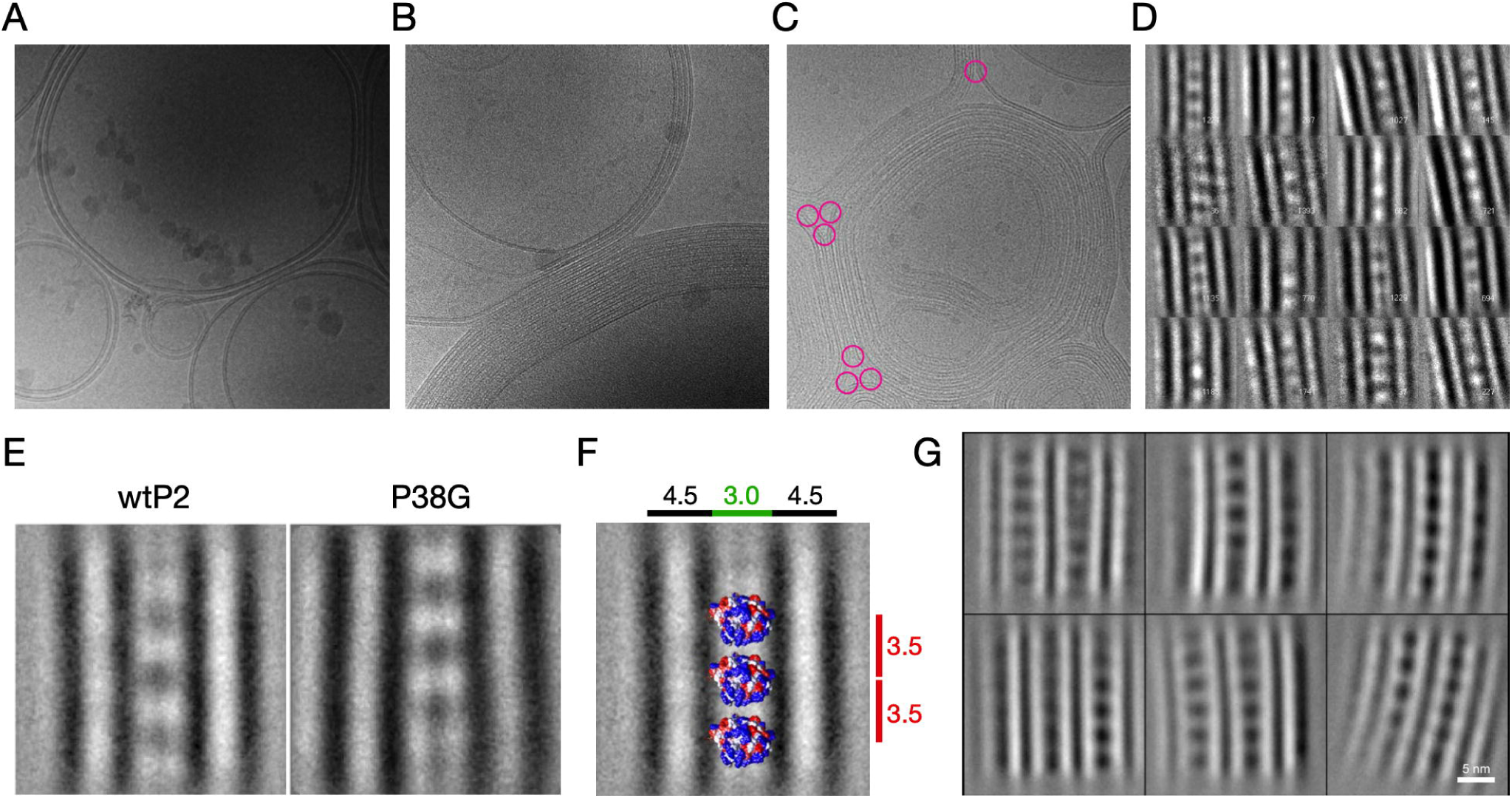
Electron cryomicroscopic analysis of lipid membrane stacking by P2. A. Cryo-EM image of *E. coli* polar lipid liposomes without protein. The images in A and B are 480×480 nm in size. B. The same liposomes in the presence of human P2 make myelin-like multilayered stacks. Lipid-to-protein mass ratio is 2.0. C. The angle between stacked membranes at the edges (pink circles) is nearly constant at ∼60°. D. 2D class averages of a single P2-linked bilayer stack. Lipid headgroups and proteins are black. The size of the box is 140×140 pixels (18×18 nm). E. Averaged structures of stacked membranes with wild-type and P38G mutant human P2. F. The space between two membranes is enough to fit one layer of P2. The membrane diameter is 4.5 nm, the space between membranes 3.0 nm, and the distance between individual P2 molecules 3.5 nm. The crystal structure of a P2 monomer has been fitted into the assembly. G. Averaging of P2-stacked multilayers, including two layers in the analysis, indicates the lattice-like arrangement of P2 throughout the myelin-like multilayer. Lipid headgroups and proteins are white; scale bar, 5 nm.

Cryo-EM was similarly carried out with the “hyperactive” P38G variant ^46^ mixed with lipids (Figure 1E). Neither the bilayer spacing nor protein-lipid organization altered in the presence of the mutant. During sample preparation, P38G induced membrane aggregation/stacking faster than wtP2, and the turbidity effect was visible within 2-3 min (not shown), in line with earlier experiments ^46^.

### 3-dimensional order in P2-bicelle complexes

In order to obtain additional structural insight into P2-membrane complexes, P2 was studied in a bicelle environment. P2 induced turbidity in protein-bicelle suspensions, and EM imaging revealed stacked arrangements of bicelles in these samples (Figure 2A). Thus, X-ray diffraction was used to gain more information on repetitive structures. In addition to the Bragg peaks originating from membrane stacking repeats of ∼7-8 nm, additional diffraction peaks were observed (Figure 2B,C) in samples with the highest lipid and protein concentrations. The corresponding distances are close to those expected from a lattice-like setup of P2 molecules between two membranes, as observed in cryo-EM. The distances can be used to deduce a possible lateral organization of P2 molecules in the membrane plane (Figure 2D).

**Figure 2.**
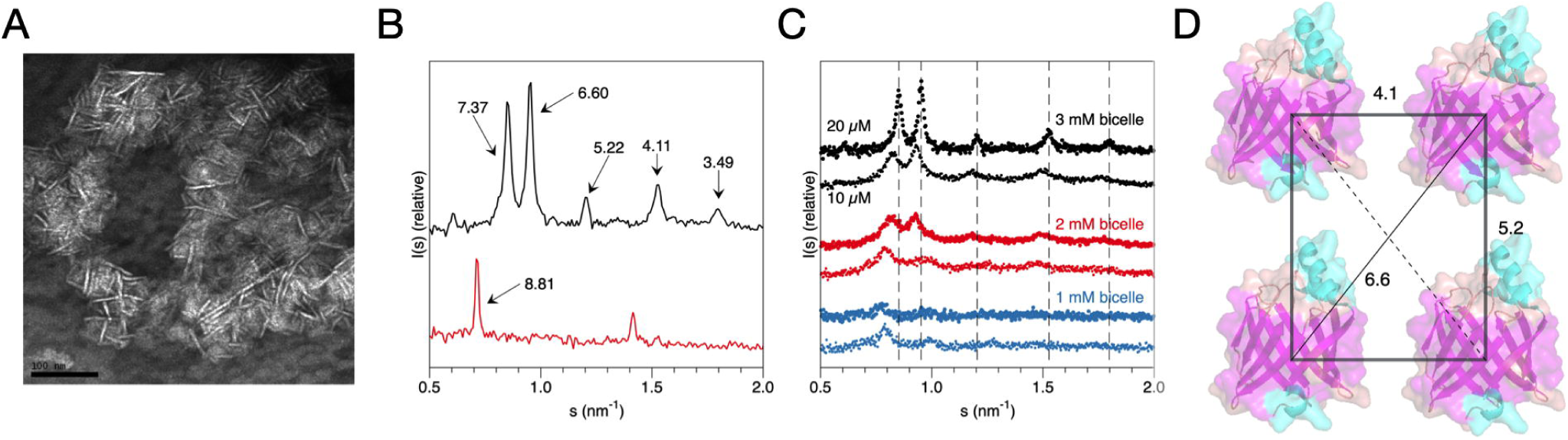
Insights into P2 structure between membranes from bicelle complexes. A. Negative staining EM micrograph of P2-stacked bicelles. Scale bar, 100 nm. B. Bragg X-ray diffraction peaks from P2-stacked bicelles (black) and vesicles (red). The corresponding repeat distances are marked. C. Titration of protein and lipid concentration in the bicelle samples indicate shorter distances and higher order when both protein and lipid concentrations increase. D. A model of P2 arrangement on the plane of the membrane, based on the peak positions in (B).

The distances observed in the experiment changed as a function of P/L ratio. This behaviour is similar, but not identical, to that observed for the P0 cytoplasmic domain, which caused tighter membrane packing at high P/L ratios ^3^. For P2, both the lipid and protein concentration affect the repeat distance in a concerted fashion (Figure 2C). The distances get shorter when lipid concentration increases, indicating an overall increase in order and tighter packing. On the other hand, at the same lipid concentration, shorter distances are observed with higher protein amounts. Hence, the protein and lipid components synergistically assemble into a compact, ordered, 3-dimensional proteolipid structure.

### Design of point mutants

In order to elucidate structure-function relationships in P2, as a general model for a FABP with a collisional mechanism and tight interaction with membranes, we used the crystal structure of human P2 to design mutations that might affect membrane binding (Figure 3A). The electrostatic surface of wtP2 shows two positively charged faces, at the helical lid domain and the bottom of the barrel structure (Figure 3B). The mutations can be roughly divided into three classes: those removing positive surface charge, those affecting the hydrophobic surface of helix α2, and other mutations possibly affecting the portal region.

**Figure 3.**
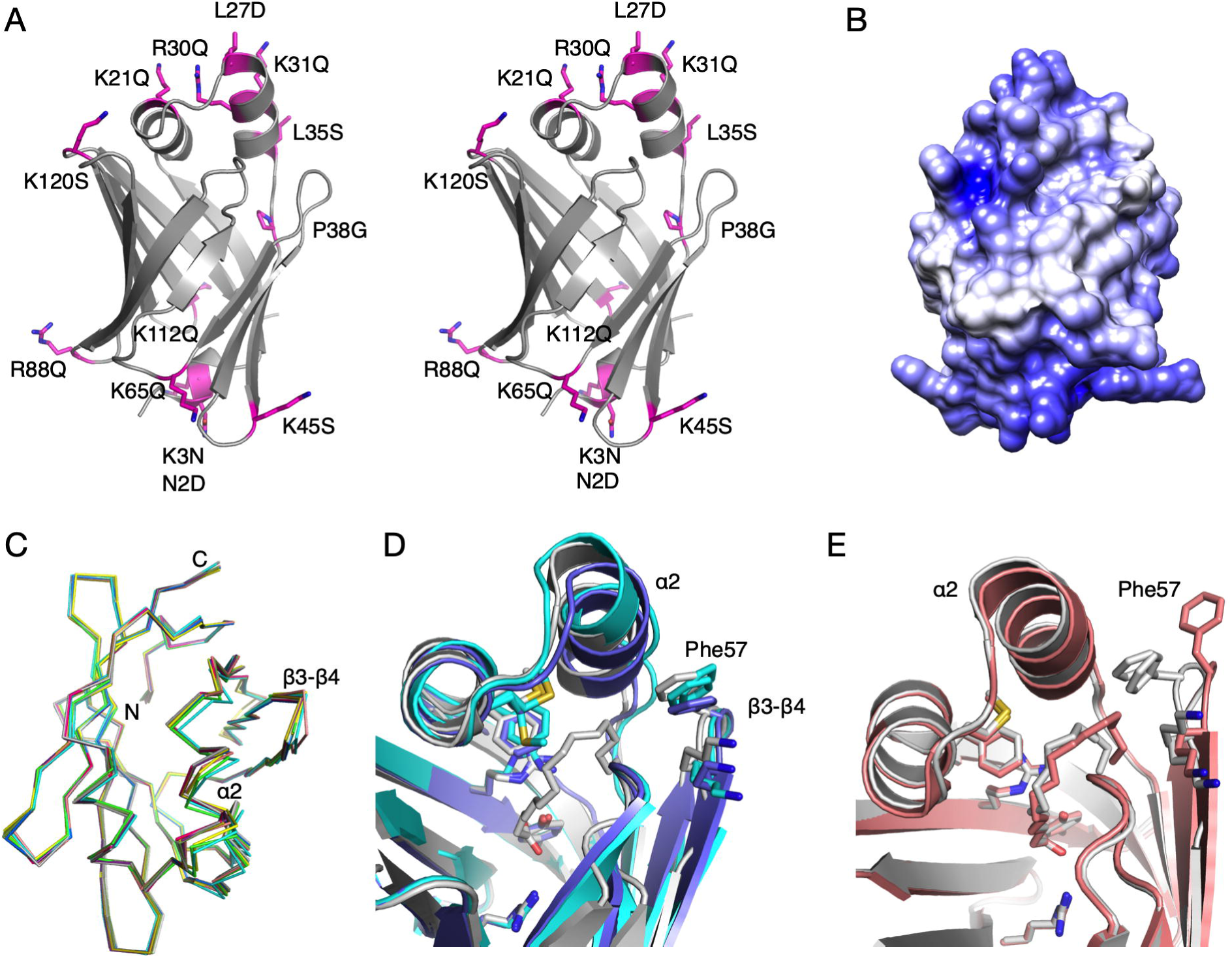
Crystal structure analysis of selected P2 mutant variants. A. Stereo view of all mutants analyzed. B. Surface electrostatics of human P2 reveal two positively charged faces at opposite ends of the molecule. C. View from the top on the C⍰ traces of all P2 variant crystal structures indicates flexibility of helix ⍰2 and the β3-β4 loop. D. Conformational differences between liganded and unliganded P38G. P38G with palmitate (PDB entry 4D6B ^46^) (grey) is superimposed with the two monomers of unliganded P38G (light and dark blue). E. Partial opening of the portal in the K65Q variant (pink), superimposed on the wtP2 structure (PDB entry 4BVM ^15^).

### Crystal structures of P2 variants

For a high-resolution insight into the structure-function differences in the P2 variants, their respective crystal structures were solved (Supplementary Table 1, Figure 3C). None of the mutations affected folding or secondary structure elements in the crystal state. With respect to this observation, it is important to note that the three studied CMT disease variants of P2 crystallized like wtP2, even though their stability and function were impaired ^30^. The RMS deviations of the mutant structures compared to wtP2 vary between 0.08 and 0.36 Å, P38G being the most divergent.

Prior to the current work, all crystal structures for wtP2 or mutant P2 contained a bound ligand inside the β barrel. The P38G structure refined here is the first exception: its internal cavity is clearly empty; no electron density for a bound fatty acid is present. This allows comparing details between liganded and unliganded P2 (Figure 3D). In our earlier study, the P38G mutant contained bound palmitate ^46^. In the unliganded crystal structure of P38G, the amino acid side chains pointing inwards mainly retain their conformation. The main-chain hydrogen bond between residue 38 and Leu10 also exists in both P38G structures. However, helix α2 at the portal region has slightly shifted outwards from helix α1 in both chains; a similar change is observed in the R30Q mutant, which could be linked to altered membrane-binding properties (see below). In addition, P38G electron density is poor for residues 33-37 at the end of helix α2 of chain B, supporting an increased flexibility/partial unfolding of the portal region in the P38G mutant in the absence of bound ligand, as seen in earlier simulations ^46^. Phe57, as well as the whole β4-β5 loop of chain A, has somewhat tilted away from the α2 helix.

All P2 structures excluding P38G have a fatty acid, modelled as a mixture of palmitate or *cis*-vaccinate in the atomic-resolution structures ^15,59^, bound inside the β barrel. The conformation and position of the fatty acid is similar in most structures. In the K65Q mutant, the conformation of the palmitate is different, and Phe57 points outwards in all four chains in the asymmetric unit (Figure 3E). This supports the proposed role for Phe57 as a gatekeeper residue in the FABP family ^13,60^.

All human P2 crystal structures published thus far have an anionic group bound in proximity of the hinge region; the identity of the ligand depends on crystallization conditions and crystal contacts. In wtP2, either chloride or citrate interacts with Thr56 and Lys37 ^14,15^. P38G and F57A contain chloride and sulfate, respectively ^13,46^. In the CMT-associated P2 mutant structures, there is a malate located in the anion binding site ^30^. In line with these data, all P2 mutant structures solved here have an anionic group bound in the same pocket. These observations lend further support to the hypothesis that this pocket may be involved in recognizing phospholipid headgroups and initiating membrane binding and/or conformational change ^15^.

### Membrane binding and multilayer stacking

SPR was used to follow binding of the P2 variants onto immobilized lipid membranes, made of either DMPC or DMPA (Figure 4A). These membranes are net neutral and negatively charged, respectively. While MBP essentially binds to lipids irreversibly on SPR ^2^, P2 dissociates from the membrane rapidly ^15^, suggesting different membrane interaction kinetics for the two proteins with overlapping function.

**Figure 4.**
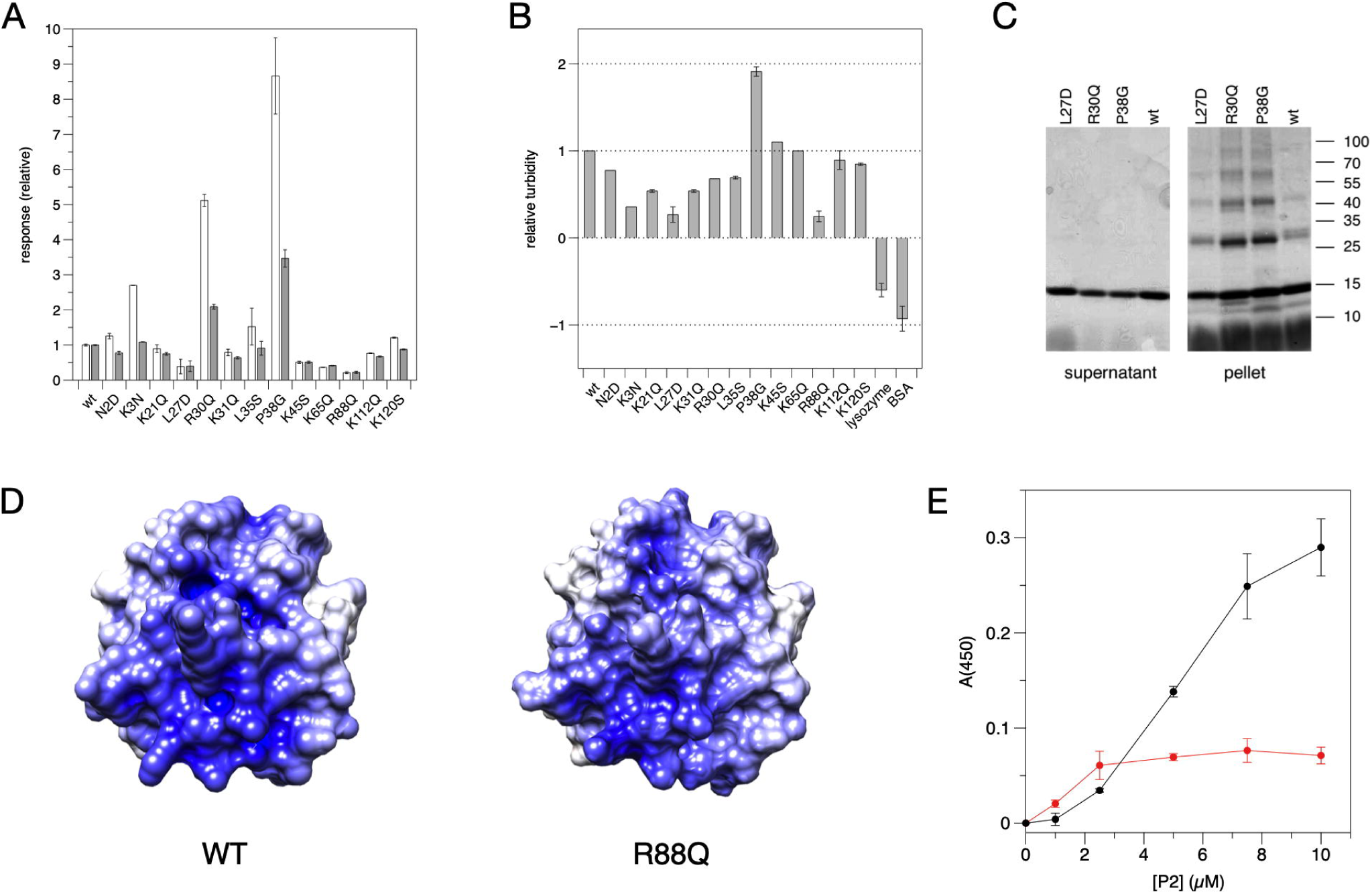
Assays on P2 variant conformation and function. A. DMPA (grey) and DMPC (white) membrane binding assays by SPR. B. Turbidity assay with 1:1 DMPC:DMPG vesicles. C. SDS-PAGE analysis of proteolipid pellets reveals SDS-resistant P2 multimers. D. Surface electrostatics of the P2 bottom surface in wtP2 (left) and the R88Q mutant (right). E. Turbidity assay of wtP2 with DMPC:DMPG vesicles (red) and bicelles (black).

Four P2 mutants showed decreased binding to lipid membranes. One of these is L27D, which affects Leu27 at the tip of the helical lid and reduces the hydrophobicity of the portal region. The other three mutations with reduced binding affinity towards lipid membranes are found in adjacent loops on the opposite face, at the bottom of the β barrel. All these mutations (K45S, K65Q, and R88Q) affect surface residues and reduce the positive charge at the bottom of the β barrel. The locations of these mutations suggest two membrane binding surfaces on opposite faces of P2, in line with its packing between two bilayers *in vivo* and *in vitro*.

While some mutations caused diminished binding to the membrane surface, P38G and R30Q had increased levels of binding. These two mutations are located in the vicinity of the portal region and the helical lid domain. The difference in membrane binding of R30Q and P38G compared to wtP2 was more pronounced, when a DMPC membrane was studied.

Turbidimetry was used to assess the effectivity of P2 variants in aggregating DMPC:DMPG vesicles (Figure 4B). When the turbid proteolipid suspensions were centrifuged and analyzed by SDS-PAGE, P2 co-sedimented with aggregated vesicles, and the strong proteolipid complex was only partially solubilized by SDS; P2 was present as a ladder of oligomeric forms (Figure 4C). Again, P38G was the most effective variant, stacking vesicle membranes more than wtP2. Some mutations caused diminished turbidity compared to wtP2. The clearest of these were L27D and R88Q; the latter lies in the β6-β7 loop - in the middle of a large positively charged surface patch at the bottom of the β barrel (Figure 4D). Hence, again, residues important for both membrane binding and stacking can be found on both positively charged faces of P2.

Another turbidimetric experiment was carried out to compare bicelles and vesicles in wtP2-induced multilayer formation. Like vesicles, bicelles are stacked by P2 into large structures causing turbidity (Figure 4E). Such ordered complexes could be a step towards higher-resolution structure determination of myelin proteolipid complexes, due to *e.g.* restrained particle size and geometry.

### Fatty acid and cholesterol binding

Using the fluorescent fatty acid analogue DAUDA, we followed internal ligand binding to P2 variants (Figure 5A). The situation is complicated by the fact that tightly bound fatty acids co-purify with P2 from the expression host. Thus, a quantitative analysis was not performed, as increased binding could reflect either higher affinity or lower amounts of copurified ligand. However, the level of bound DAUDA should correlate with the opening of the portal region and/or the barrel, which is also required for removal of the bound fatty acid. Since bound fatty acid affects dynamics of P2 ^13,46^, it is likely that some of the mutated variants have different affinities towards fatty acids. Most mutant variants showed slightly higher DAUDA signal than wtP2, and P38G was the strongest binder of all variants.

**Figure 5.**
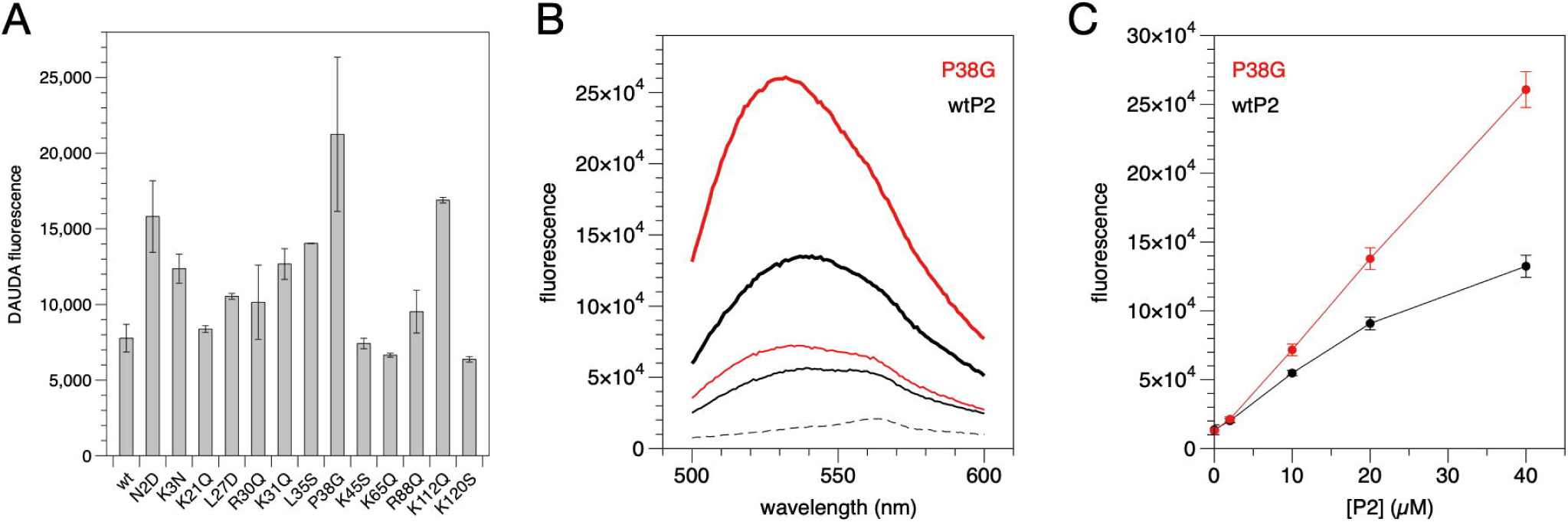
Ligand binding by human P2. A. Binding of the fluorescent fatty acid DAUDA. B. Binding of NBD-cholesterol by wtP2 (black) and P38G (red). Dashed line, ligand alone; thin line, 10 µM P2; thick line, 40 µM P2. C. Concentration dependence of fluorescence at 532 nm.

We previously proposed cholesterol binding by P2 ^14^, in addition to fatty acids. Cholesterol binding was tested using wtP2 and P38G. wtP2 induced a clear, concentration-dependent change in the fluorescence spectrum of the environment-sensitive probe 22-NBD-cholesterol; the fluorescence maximum shifted towards shorter wavelengths and its intensity increased (Figure 5B,C). The spectral changes were more pronounced with P38G, which has a more flexible portal region ^46^. The experiment shows that cholesterol, which is very abundant in myelin, could be a physiologically relevant ligand for P2. These assays together indicate that protein flexibility is important when P2 binds to its biological ligands.

### Folding and stability of point mutant variants

CD spectroscopy was used to analyze the folding and stability of the P2 variants. While most mutations had little effect on thermal stability, P38G had two steps of unfolding, the first one appearing already at 50 °C and the second one only at >75 °C (Figure 6A). Another outlier was R30Q, which had a slightly lowered stability compared to wtP2. Interestingly, these two mutations causing changes in stability are those with enhanced membrane binding and stacking properties. In the crystal structures, they present minor conformational differences in their helical lid, compared to wtP2.

**Figure 6.**
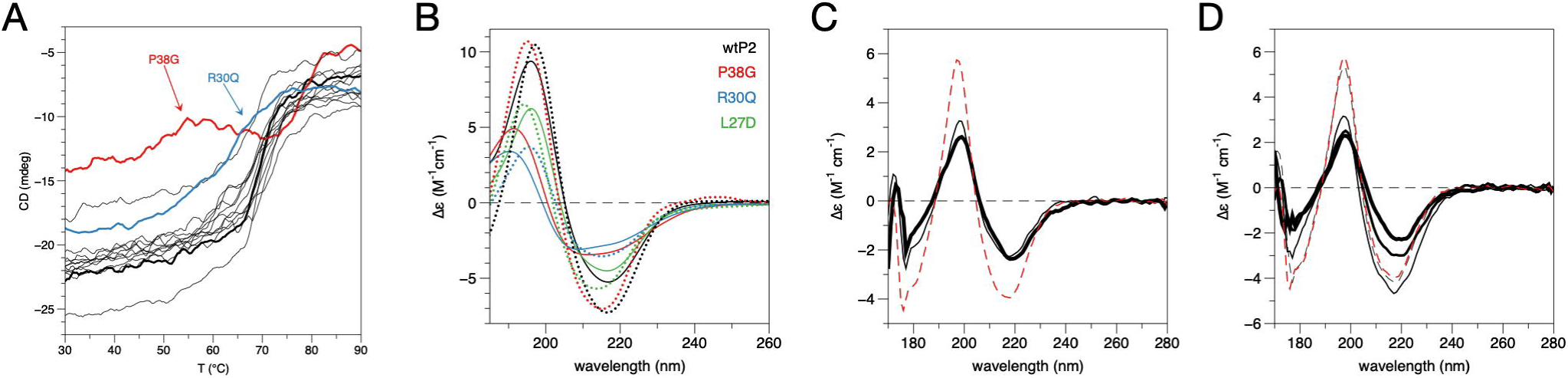
P2 stability and folding. A. Melting curves for wtP2 and all studied mutants. The outliers are P38G (red) and R30Q (blue). The thick black line represents wtP2. B. Conformation of wtP2 and selected mutants in the presence (solid lines) and absence (dotted lines) of DMPC:DPC bicelles. C. Wild-type P2 in water (red dashed line), 9:1 DMPC:DMPG (thin black line), and 1:1 DMPC:DMPG (thick black line). D. Wild-type P2 in water (red dashed line), and lipid:DPC bicelles containing DMPC (black dashed line), 9:1 DMPC:DMPG (thin black line), 4:1 DMPC:DMPG (medium black line), and 1:1 DMPC:DMPG (thick black line).

To elucidate the conformational changes induced by membrane binding, SRCD spectra for wtP2 and some divergent mutants were measured in the presence and absence of DMPC:DPC bicelles (Figure 6B). For wtP2, bicelle binding induced small changes in the SRCD spectrum. L27D exhibited less change in the spectrum in the presence of bicelles, supporting the reduced membrane binding of L27D observed in SPR and turbidity assays. On the other hand, P38G, having a higher propensity for membrane interactions, showed larger conformational changes in the bicelle environment. R30Q behaved much like P38G; both variants showed partial unfolding. These results highlight the importance of protein flexibility in membrane binding and indicate a role for the α-helical lid in P2-membrane interactions.

The bicelle system was used to deduce effects of lipid composition on wtP2 folding state, as well as to compare to vesicles with the same lipid composition. SRCD spectra showed that in vesicles, both 1:1 and 9:1 DMPC:DMPG gave the same conformational change of wtP2 compared to the protein in water (Figure 6C). In bicelles, however, wtP2 behaved differently, in that very little change occurred in DMPC alone or in 9:1 DMPC:DMPG, and the spectrum changed more with 1:1 and 4:1 DMPC:DMPG, to resemble the one measured with vesicles. (Figure 6D). These differences with respect to lipid composition could be related to membrane curvature.

### Atomistic simulations on P2-membrane interactions

To combine aspects of high-resolution structural data and membrane binding, we studied the interactions of wild-type and P38G P2 with membrane surfaces using atomistic MD simulations (Figure 7). Two membrane systems were built: a 1:1 mixture of DMPC:DMPG, which corresponds to compositions often used in the laboratory, and a myelin-like membrane based on literature values ^49^.

**Figure 7.**
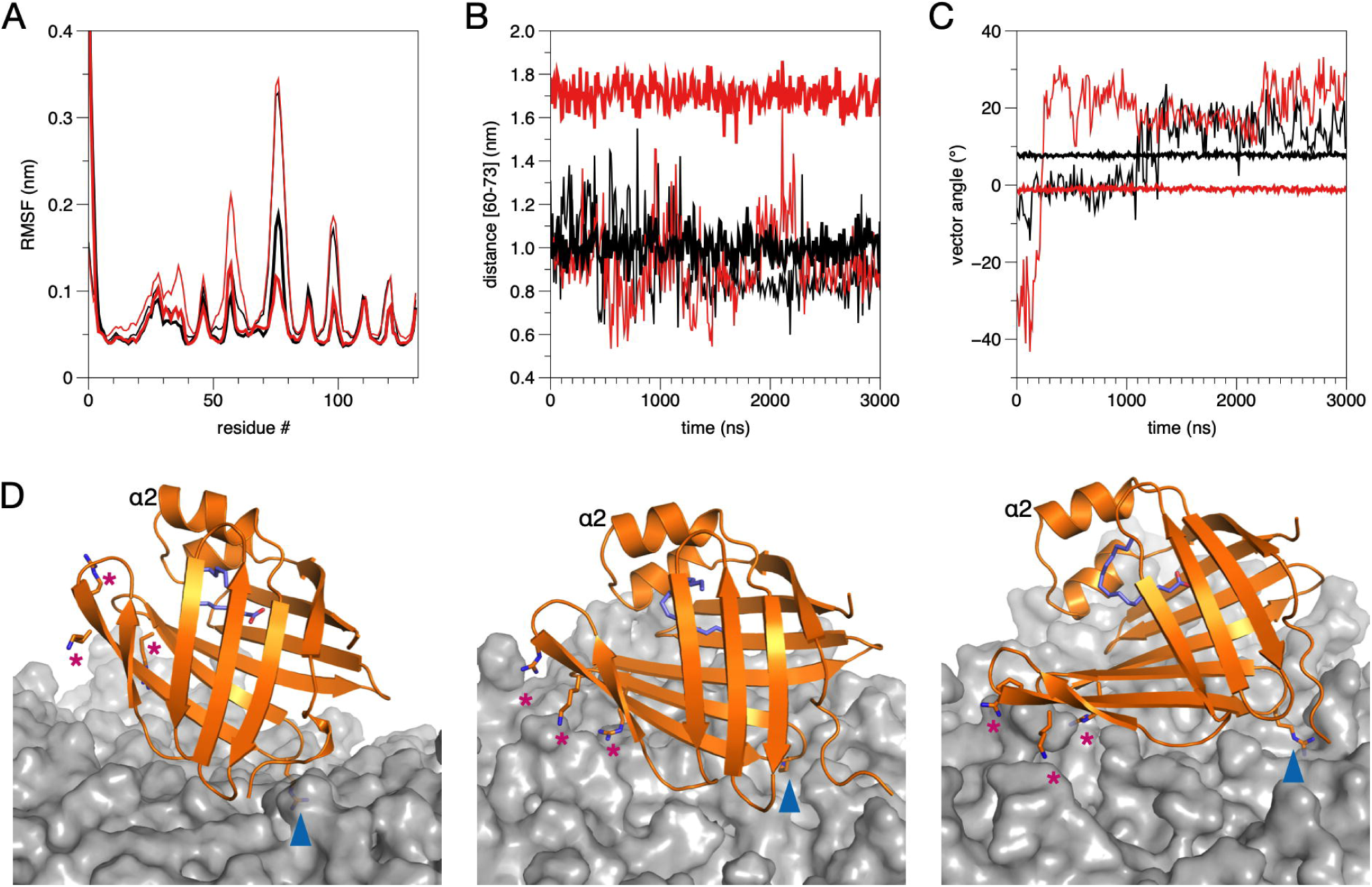
MD simulations on P2 binding to lipid membrane surface. A. RMSF for wtP2 (black) and P38G (red) in 1:1 DMPC:DMPG (thin lines) and myelin lipid composition (thick lines). B. Distance of the β4-β5 opening of the β barrel during the simulation. Colouring as in (A). C. Angle of the P2 β barrel axis with respect to the membrane surface. Note how both wtP2 and P38G are rigidly anchored to the same orientation immediately after the equilibration period. D. Snapshots from the simulations. Left: wtP2 on DMPC:DMPG at 1135 ns. Middle: wtP2 on myelin at 2060 ns. Right: P38G on myelin at 800 ns. Locations of the two membrane anchors, Arg88 (blue arrowhead) and Arg78/Lys79/Arg96 (magenta asterisks) are indicated.

During the attachment of wtP2 onto the membrane surface, a similar orientation was always observed: this involved the positively charged surface close to the bottom of the β barrel. Arg88 is a central residue in initial P2-membrane interactions. While it was expected that initial membrane binding would involve the portal region and the helical lid, this orientation, with the bottom face of the β barrel first approaching the membrane, is reproducible. The protein was further turned on its side in this arrangement in the myelin lipid composition, indicating that the rows of P2 molecules observed in cryo-EM images do not embed deep into the bilayers. The 3-nm spacing between membranes can accommodate one layer of P2 in this orientation.

A difference in orientation was observed between the DMPC:DMPG and myelin membranes; P2 remains more upright and dynamic in DMPC:DMPG, while it falls rigidly on its side on the myelin-like membrane (Figure 7C). Differences in wtP2 dynamics were additionally observed between lipid compositions. The protein was more rigid when bound to the myelin membrane (Figure 7A); on DMPC:DMPG, it had higher dynamics and hung on the membrane with the Arg88 anchor (Figure 7C). During the simulation with the myelin-like bilayer, the PIP_2_ molecules within the myelin bilayer bound to the tip of the β5-β6 and β7-β8 loops, promoting opening of the β barrel, while Arg88 at the other end of the protein, in the β6-β7 loop, interacted strongly with POPS head groups (Supplementary Figure 1, Figure 7D). The PIP_2_ binding site is formed of the side chains of Arg78, Lys79, and Arg96. These results could reflect an important difference between a biological membrane composition and simplistic membrane models.

The electrostatic interactions of wtP2 and P38G were very similar with the membrane lipids during the simulations (Supplementary Figure 1). The P38G variant similarly attached to the myelin-like membrane surface, being anchored sideways, and opened up even more than wtP2 (Figure 7B,D). The portal region, and the expected opening during ligand exchange ^13^, face upwards in this setting, and upon the approach of another membrane, they could closely interact with its surface.

## Discussion

Myelin protein P2 is a unique member of the FABP family, able to stack lipid bilayers together, in addition to being a member of the FABP subgroup carrying out collisional transfer. Lipid membrane binding by P2 involves the hydrophobic tip of the helical lid, electrostatic interactions, and dynamics of the portal region ^13,15,46^. Here, we have revealed details of the assembly of the P2-membrane stacks and the surprising role of the bottom region of the P2 β barrel in membrane binding. The data provide much-needed information on the assembly of the myelin membrane at the molecular level.

### Structure of P2-induced proteolipid multilayers

Our cryo-EM experiments illustrate an organized lattice-like supramolecular 3-dimensional arrangement of P2-membrane stacks. Surprisingly, P2, a 15-kDa protein, which has dimensions of 4.5 nm x 3.6 nm, is visible between the lipid bilayers as a lateral network. Both the cryo-EM images and calculated 2D class averages of P2-membrane stacks show a constant distance (3 nm) between the apposing lipid membranes and a repeat distance (containing a single bilayer and intermembrane space) of 7.5 nm. Earlier, a repeat distance of ∼9 nm for P2-membrane stacks was measured by X-ray diffraction using DMPC:DMPG in suspension ^15^. Bragg peaks in X-ray diffraction experiments support the highly organized arrangement of P2-membrane stacks seen in cryo-EM, and the conditions for preparing cryo-EM samples, with less hydration, might be more relevant to myelin *in vivo*. Indeed, using the bicelle model system, we measured repeat distances of 7.5 nm in stacks of bicelles induced by P2, and the distance evolved as a function of protein and lipid concentration. Thus, P2 may have a function in defining the membrane spacing in PNS compact myelin, together with MBP and P0. All three of these proteins produce membrane stacks *in vitro* ^2,3^, with intermembrane spacing very close to that seen in the mature myelin major dense line.

The spacing between the neighboring P2 molecules between membrane bilayers is constant (∼3.5 nm), and there appears to be a relationship between the positioning of P2 molecules between consecutive membrane layers. The results suggest the presence of a near-crystalline lattice of P2 between membranes; this is also supported by our X-ray diffraction experiment using stacked bicelles, in which - unlike earlier similar experiments using lipid vesicles - we see new distances much shorter than those coming from bilayer stacking *per se*. As these distances depend on protein concentration, they correspond to distances between proteins arranged as a lateral layer between two membranes. Whether such packing occurs *in vivo*, depends on the local P2 concentration in myelin as well as the presence and organization of other highly abundant myelin proteins, such as P0 and MBP. The quantity of P2 has been reported to vary between different regions of PNS as well as from nerve fiber to nerve fiber ^12^.

We recently reported, using similar cryo-EM approaches, the arrangement of the extracellular domains of P0 as a zipper-like assembly between the membranes ^3^. The assembly of P2 at the cytoplasmic face shown here completes the picture of PNS myelin molecular assembly. Importantly, while P0 extracellular domains interact with each other as two layers between membranes, only a single layer of P2 is observed, and each protein molecule must interact with two cytoplasmic leaflets simultaneously. The details of this aspect were further characterized here through mutagenesis, functional experiments, and high-end computer simulations.

### Functional residues revealed by point mutations

The unique ability of P2 to stack lipid membranes requires two membrane-binding sites on opposite faces of the protein; P2 has two positively charged surfaces. Membrane binding experiments for surface-mutated P2 gave information about crucial regions and mechanisms of protein-membrane interaction. The L27D mutation at the tip of the α-helical portal region reduces membrane stacking and binding, as well as diminishes the changes in CD spectrum upon introducing membrane-mimetic bicelles. Thus, Leu27 may be inserted into the hydrophobic core of a lipid bilayer. This insertion is presumably facilitated by a conformational change in the portal region ^15^. In addition, other portal region mutations (K21Q, K31Q), which remove a positive charge, also decreased membrane binding and stacking. These residues probably interact with negatively charged lipid head groups and, together with Leu27, form a membrane anchor of the P2 portal region. We earlier showed that the L27D mutation impairs the formation of stacked membrane systems in a cell culture system ^15^.

On the other hand, the removal of a positive charge at the opposite end of the β barrel (mutations K45Q, K65S, R88Q and K112Q) caused reduced membrane binding and stacking. In MD simulations, R88Q protrudes into the lipid membrane and forms tight interactions with lipid head groups, especially PS in the myelin-like bilayer. However, there are no hydrophobic residues at the bottom face of P2, and the barrel bottom interaction with the lipid membrane is facilitated by electrostatic interactions. The bottom region of P2 is unlikely to be deeply inserted into membranes, nor will it undergo large conformational changes.

An exception within all P2 mutants concerns P38G. In line with earlier data ^61^, it is more active in most of the experiments, including membrane stacking as well as membrane, fatty acid, and cholesterol binding assays. The P38G mutation, however, does not alter the organization or repeat distance of the P2-membrane stacks in cryo-EM. In the crystal structure of P38G, there is no fatty acid bound, and this mutant is more flexible and has altered dynamics compered to wtP2 ^61^. In MD simulations, the fatty acid was observed nearly escaping from the barrel ^13^. The weak electron density of the portal region in the P38G mutant crystal structure supports the idea of a flexible lid in this mutant, making it more dynamic ^46,62^ and prone to opening. The other mutation, R30Q, which increased the flexibility of the portal region, causes smaller but similar effects on the activity of P2 in several assays, confirming the importance of the dynamics of the portal region in the function of P2. For other FABPs, the Arg residue corresponding to P2 Arg30 has been suggested to attract negatively charged fatty acids ^63,64^; while this could be happening in P2 as well, the R30Q mutation clearly has larger-scale effects on membrane interactions and local folding or dynamics.

Phe57 is a conserved residue within the FABP family suggested to be a general gatekeeper for ligand binding ^13,60^. It controls ligand entry into the β barrel and can flip between two conformations ^13,65^. Phe57 points outwards in the K65Q crystal structure, and the fatty acid shifts towards the opening cleft. It is unclear how a mutation located at the other end of the β strand might induce the flipping of Phe57. However, the Phe57 flip may be one initial step in P2 opening, which was observed in MD simulations and structural studies ^13,30^.

Arg88 appears central to the initial P2-membrane contact, functioning as an anchor. Within the human FABP family, Arg88 is conserved in P2, but not other family members ^15^. This indicates its possible importance for the membrane stacking function, since other collision-type FABPs, bind transiently to single membrane surfaces.

### Conformational changes and dynamics upon membrane binding

The binding of P2 onto a myelin-like membrane is accompanied by a conformational change opening the likely entry/egress site of the bound fatty acid. This change can be observed both experimentally and in computer simulations. The change is similar to that observed in solution for the CMT disease variants and in extended computer simulations of P2 ^13,30,46^. Similar conformational changes were observed for H-FABP during long simulations ^63^. The bound ligand could be exchanged with the apposing membrane in a multilayer, when this conformational change occurs.

Atomistic simulations of P2 on a membrane surface revealed different behaviour on a simplistic model membrane compared to a myelin composition. Importantly, some of the lipids concentrated on the myelin membrane formed specific interactions with P2 during the long atomistic simulations, contributing to the conformational change. The presence of two anchor points for P2 membrane binding enabled the membrane to contribute to P2 barrel opening, unraveling the bound ligand. The negatively charged lipids, PS and PIP_2_, might also affect other myelin proteins in a specific fashion, and further experiments will be required to grasp the full scope of intertwined interactions between myelin proteins and specific lipids. A model combining current data on P2 bound to the cytoplasmic leaflet of myelin is shown in Figure 8.

**Figure 8.**
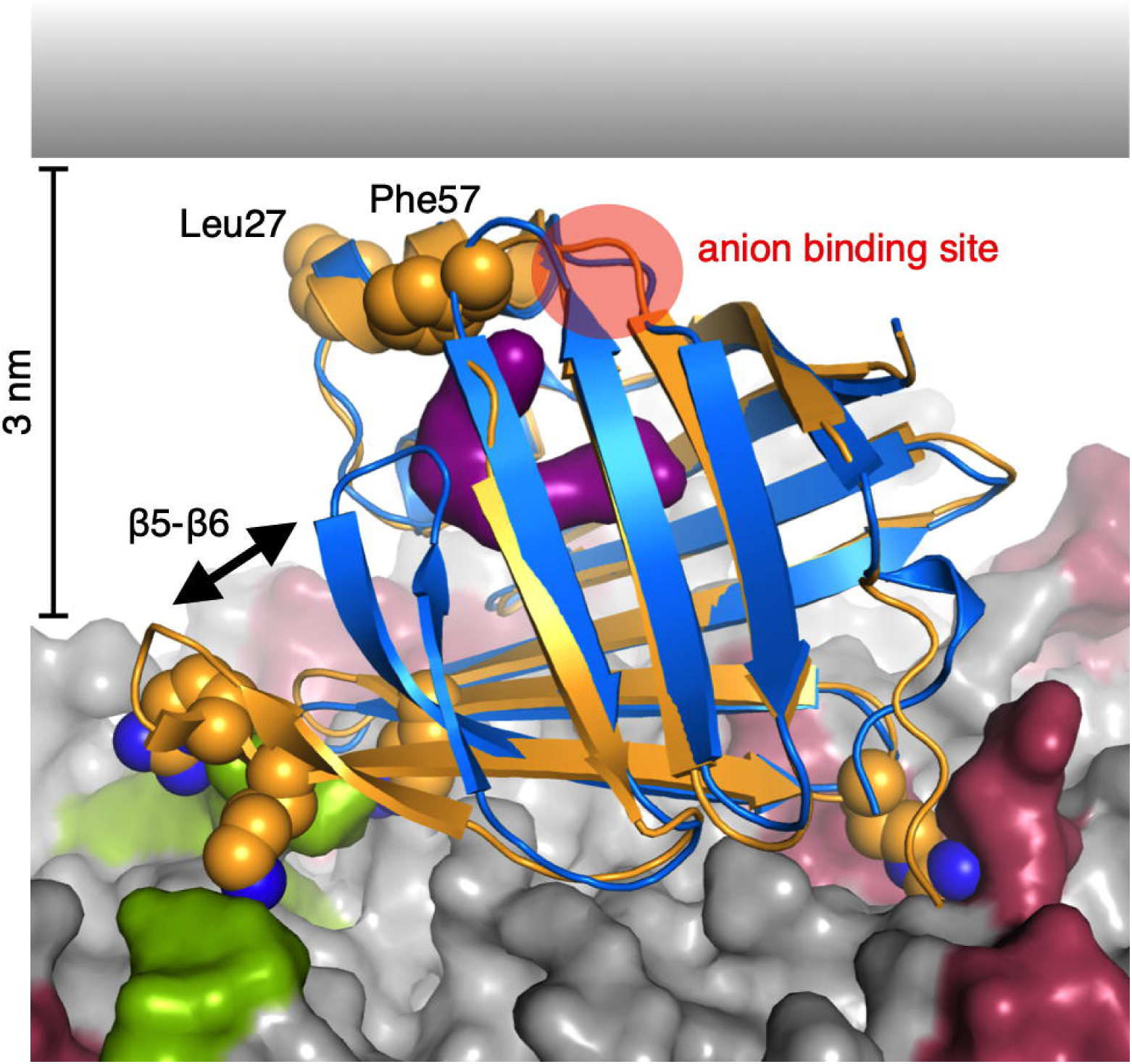
Model for P2-membrane interactions based on current data. Shown is a superposition of wtP2 crystal structure (blue) and the membrane-bound conformation of P38G (orange). Upon membrane binding, Arg88 gets anchored by POPS molecules (red) and the basic residues around the β5-β6 loop interact strongly with PIP_2_ (green). The opening of the β5-β6 flap exposes the fatty acid ligand (purple). Leu27, Phe57, and the anion binding site are facing the apposing membrane surface in this setting.

Upon the formation of a P2-membrane complex, the dynamics of both the protein and lipid components are altered. When bound to P2, the dynamics of the lipid membrane are decreased ^56^, while P2 becomes extremely heat-stable when bound to membranes ^15^. These observations are likely to be linked to the synergistic effects of P2 and the lipids in the tightening of the 3D molecular assembly, as shown by X-ray diffraction from myelin-mimicking bicelle complexes. Furthermore, they are in line with the decreased dynamics of P2 on a myelin-like membrane in the simulations.

## Concluding remarks

We have shown that, similarly to P0, MBP, and PMP22, myelin protein P2 is able to spontaneously induce the formation of myelin-like membrane multilayers. We have for the first time visualized the arrangement of P2 between membranes, providing an unprecedented view into the structure of the major dense line in peripheral nerves. Furthermore, our observations provide a lipid composition-dependent mechanism for the opening of the P2 structure for ligand entry and egress; in the case of a multilayered membrane, the ligand could be exchanged with the apposing membrane. How myelin proteins act together in forming native myelin multilayers through interactions at both extracellular and intracellular surfaces of the bilayer is a major question in myelin biology; the tools and materials exist for solving this question in the coming years.

## Acknowledgements

The authors wish to thank financial support granted by the Academy of Finland, the Emil Aaltonen Foundation (Finland), Jane and Aatos Erkko Foundation (Finland), the Helsinki Institute of Life Science Fellow programme (Finland), the Science and Research Foundation of the City of Hamburg (Germany), and the Sigrid Jusélius Foundation (Finland). Beamtime and user support at EMBL/DESY and ISA are gratefully acknowledged. Travel to synchrotrons was supported by the Norwegian Research Council (SYNKNØYT project 247669) and EU Horizon 2020 programmes iNEXT (grant 653706) and CALIPSOplus (grant 730872).

## Author contributions

S.R., M.L., I.V., H.S., and P.K. conceived of the study. S.R., O.C.K., J.K., T.N., M.L., A.R., V.P.D., and P.K. carried out the experiments. All authors contributed to the interpretation of the results. S.R. and P.K. took the lead in writing the manuscript. All authors provided critical feedback and helped shape the research, analysis, and manuscript.

## Figure legends

**Supplementary Figure 1. Electrostatic interactions between key basic residues and membrane lipids.** Green, PIP_2_; black, POPS; red, cholesterol; blue, POPE; magenta, POPC; orange, sphingomyelin. Note how Arg78, Lys79, and Arg96 have stable PIP_2_ contacts, while Arg88 has strong interactions with PS, and these interactions are observed for both wtP2 and P38G.

