## Supplementary Table 1 for "Determinants for forming a supramolecular myelin-like proteolipid lattice"

### Supplementary Table 1. Crystallographic data processing and refinement.

| mutant | N2D | K3N | K21Q | L27D | R30Q | K31Q | L35S | P38G | K45S | K65Q | R88Q | K112Q | K120S |
|---|---|---|---|---|---|---|---|---|---|---|---|---|---|
| space group | $P4_12_12$ | $P4_12_12$ | $P4_12_12$ | $P4_12_12$ | $P2_12_12_1$ | $P4_12_12$ | $P4_12_12$ | $P2_12_12_1$ | C2 | $P4_3$ | $P4_12_12$ | $P4_12_12$ | $P4_12_12$ |
| unit cell dimensions (Å,Å,Å, °,°,°) | 58.26 58.26 101.72 90 90 90 | 64.5 64.5 101.2 90 90 90 | 57.96 57.96 101.24 90 90 90 | 64.88 64.88 100.92 90 90 90 | 58.59 75.96 100.63 90 90 90 | 58.24 58.24 102.19 90 90 90 | 65.83 65.83 101.32 90 90 90 | 55.54 65.49 82.23 90 90 90 | 120.13 63.61 94.47 90 130.1 90 | 83.03 83.03 77.90 90 90 90 | 64.88 64.88 264.35 90 90 90 | 65.71 65.71 101.19 90 90 90 | 63.68 63.68 101.4 90 90 90 |
| resolution range (Å) | 30-1.65 (1.70-1.65) | 20-2-70 (2.77-2.70) | 30-2.30 (2.36-2.30) | 30-2.31 (2.37-2.31) | 25-3.0 (3.08-3.00) | 40-1.80 (1.85-1.80) | 25-2.00 (2.05-2.00) | 20-1.80 (1.85-1.80) | 20-2.20 (2.26-2.20) | 20-1.20 (1.23-1.20) | 40-1.80 (1.85-1.80) | 25-1.80 (1.85-1.80) | 40-2.90 (2.98-2.90) |
| completeness (%) | 97.4 (78.9) | 99.5 (100) | 99.2 (97.6) | 98.9 (94.6) | 98.4 (95.5) | 98.7 (99.8) | 98.2 (99.8) | 90.9 (58.4) | 88.0 (49.2) | 93.9 (63.2) | 99.4 (96.8) | 99.4 (95.6) | 99.6 (97.4) |
| <I/sigmaI> | 15.2 (1.5) | 14.5 (3.1) | 11.1 (2.2) | 8.0 (0.9) | 9.0 (1.2) | 19.0 (2.5) | 19.8 (1.7) | 21.8 (2.2) | 11.0 (1.6) | 19.5 (1.4) | 19.8 (2.0) | 30.2 (2.0) | 10.5 (1.8) |
| redundancy | 4.7 (2.0) | 10.7 (11.2) | 5.1 (4.7) | 7.9 (4.7) | 4.7 (4.6) | 9.5 (9.2) | 12.3 (9.7) | 5.5 (3.7) | 3.4 (2.8) | 5.1 (2.1) | | 11.8 (11.2) | 7.2 (7.3) |
| $R_{sym}$(%) | 8.0 (54.1) | 17.3 (80.0) | 13.8 (75.5) | 21.7 (197.1) | 12.6 (168.0) | 11.1 (92.6) | 10.0 (149.2) | 4.9 (57.6) | 7.8 (62.3) | 4.1 (66.1) | 7.9 (77.3) | 6.1 (126.7) | 13.3 (106.5) |
| completeness after anisotropic truncation | | | | 81.5 | 84.7 | | 86.3 | | | | | | |
| $R_{work}$ (%) | 14.6 | 21.2 | 18.8 | 18.8 | 25.4 | 14.5 | 17.0 | 16.7 | 19.4 | 13.1 | 15.9 | 17.8 | 21.0 |
| $R_{free}$ (%) | 18.8 | 25.0 | 22.4 | 24.5 | 28.9 | 18.9 | 21.3 | 22.5 | 25.2 | 15.4 | 19.9 | 20.3 | 25.7 |
| RMSD bond length (Å) | 0.009 | 0.002 | 0.003 | 0.003 | 0.003 | 0.008 | 0.012 | 0.012 | 0.004 | 0.010 | 0.010 | 0.012 | 0.003 |
| RMSD bond angle (°) | 1.1 | 0.5 | 0.6 | 0.5 | 0.6 | 1.0 | 1.0 | 1.2 | 0.8 | 1.1 | 1.1 | 1.4 | 0.5 |
| Ramachandran favoured (%) | 99.3 | 99.2 | 98.5 | 98.5 | 93.9 | 100 | 98.6 | 96.5 | 98.0 | 99.2 | 100 | 100 | 96.2 |
| Ramachandran outliers (%) | 0 | 0 | 0 | 0 | 0.4 | 0 | 0 | 0.4 | 0 | 0 | 0 | 0 | 0 |
| Molprobity score, percentile | 1.15, 99th | 1.31, 100th | 0.99, 100th | 0.67 (100th) | 2.03 (99th) | 1.17 (99th) | 1.14, 100th | 1.62, 89th | 1.82 (93rd) | 1.41 (81st) | 1.45 (96th) | 1.78 (80th) | 1.68 (100th) |
| PDB entry | 6xu5 | 6xu9 | 6xua | 6xuw | 6sts | 6xvq | 6xvr | 6xvs | 4a1h | 4a1y | 6xvy | 4a8z | 6xw9 |
