## Supplementary figures and images for "Determinants for forming a supramolecular myelin-like proteolipid lattice"

### Supplementary Figure 1

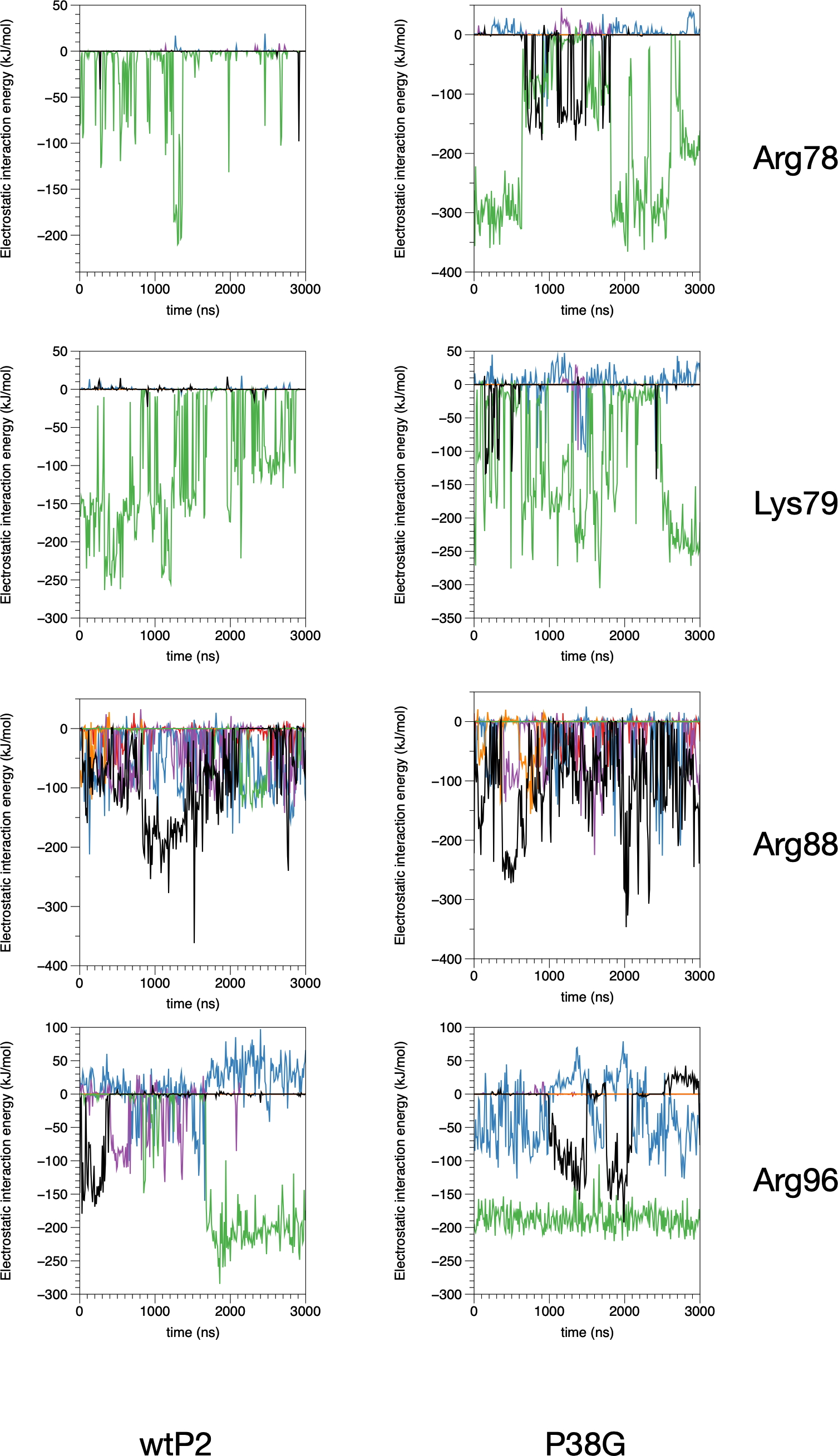
